# Increased alcohol self-administration following repeated Toll-like receptor 3 agonist treatment in male and female rats

**DOI:** 10.1101/2021.10.26.465779

**Authors:** Dennis F. Lovelock, Patrick A. Randall, Kalynn Van Voorhies, Ryan P. Vetreno, Fulton T. Crews, J. Besheer

## Abstract

Toll-like receptor (TLR) signaling may play an important role in the neuroimmune system’s involvement in the development and maintenance of alcohol use disorder. In the present study we administered TLR3 agonist poly(I:C) in male and female Long-Evans rats to determine whether TLR3 agonism can increase alcohol consumption in a daily 15% alcohol operant self-administration paradigm. We found few effects when poly(I:C) was given every-other-day at 0.3 or 1.0 mg/kg, however when instead 1.0 mg/kg was given on consecutive days alcohol intake increased in the days following injections specifically in females. Furthermore, in a second experiment we found that this effect only emerged when rats had a history of multiple poly(I:C) injections. In the final experiment the dose was increased to 3.0 mg/kg on consecutive days which resulted in significant reductions on injection days in females that were not accompanied by subsequent increases. The dose was increased to 9 mg/kg for one final pair of injections which led to reductions in intake in both males and females but only increased subsequent alcohol consumption in males. Overall, poly(I:C) was able to increase subsequent alcohol consumption in both sexes, with females being sensitive to lower doses than males both in terms of changes in alcohol consumption and general locomotor reduction. These findings show that TLR3 agonism may be involved in driving increased alcohol consumption and add to the body of work identifying the neuroimmune system as a potential therapeutic target for AUD.

## Introduction

Alcohol use disorder (AUD) is a widespread public health issue affecting millions of people each year. An area that has gained considerable attention is the role that the innate immune or neuroimmune system plays in the development and maintenance of alcohol use and abuse (Blednov et al., 2012; Crews et al., 2017a; Crews et al., 2013; Erickson et al., 2019; Mayfield et al., 2013). Studies in postmortem human brain tissue find that proinflammatory signaling molecules are increased in AUD (Coleman et al., 2017; Crews et al., 2013; Crews and Vetreno, 2014; Crews et al., 2016; Crews et al., 2017b; Qin and Crews, 2012; Qin et al., 2021). Toll-like receptors (TLRs), part of the interleukin-1/TLR superfamily responsible for generating inflammatory responses, are a crucial component of the neuroimmune system (Crews et al., 2017b). These pathogen detecting receptors comprise one of the earliest phases of the inflammatory response to infection and other agents both in the brain and periphery. In the past decade, there has been mounting evidence that these receptors also play an important role in the development of AUDs (Crews and Vetreno, 2014; Crews et al., 2017b; McCarthy et al., 2017). Indeed, chronic alcohol use has been linked to increased expression of TLRs in the brain that correlates with lifetime alcohol intake (Qin and Crews, 2012). Furthermore, activation of these receptors has been shown to increase alcohol intake (Grantham et al., 2020; Randall et al., 2020; Warden et al., 2019b). As such, alcohol-induced activation of neuroimmune signaling via TLRs likely leads to the development of feedback loops supporting continued alcohol intake.

To date, several types of TLRs have been implicated in alcohol drinking in preclinical models. TLR4 receptors, for instance, have been shown to be upregulated following binge alcohol consumption, adolescent alcohol exposure, and following treatment with endotoxin lipopolysaccharide (Blednov et al., 2011). Treatment with a TLR7 small molecule agonist has been shown to increase home cage ethanol consumption (Grantham et al., 2020). Expression of TLR3 receptors is increased following alcohol intake (Warden et al., 2019a)and treatment with the TLR3 agonist polyinosinic:polycytadylic acid (poly(I:C)) increases alcohol intake in mice and rats (Qin and Crews, 2012; Randall et al., 2019; Warden et al., 2019a, b). Furthermore, genetic knockout of TLR3 not only decreases alcohol intake but also decreases acute tolerance to alcohol in male mice further suggesting an important role of TLR3 in both alcohol intake and physiological response to alcohol (Blednov et al., 2021).

Interestingly, TLR3 is also the only TLR that does not signal through the myeloid differentiation primary response-88 (MyD88) dependent pathway, instead relying on the TIR-domain-containing adaptor inducing interferon-β-dependent (TRIF) pathway (Kawasaki and Kawai, 2014). This is an important difference as TRIF signaling has been shown to produce opposing effects to MyD88. For example, mice lacking MyD88 are less sensitive to the sedative and ataxic effects of alcohol and show increased alcohol intake (Blednov et al., 2017). By contrast, inhibiting IKKε and TBK1, downstream components of the TRIF pathway, male mice decrease alcohol intake (McCarthy et al., 2018). Together, this suggests an important regulatory role for TLRs, in particular TLR3, in regulating alcohol intake.

A previous study from our lab assessed the effects of poly(I:C) on alcohol self-administration in rats and its’ effects on glutamatergic signaling (Randall et al., 2019). Similar to others, we found that activation of TLR3 increased alcohol intake. In addition, we found that a single injection of poly(I:C) (3 mg/kg, IP) was capable of producing rapid increases in neuroimmune-related gene expression (6 hours post-injection) that remained elevated several weeks later. These findings led to the question of whether repeated dosing with poly(I:C) would further enhance alcohol intake. That is, we hypothesized that repeated injections with poly(I:C) would prime or sensitize the system (Crews and Vetreno, 2018) to further enhance later alcohol intake. To test this, the present experiments used multiple repeated poly(I:C) injection protocols in male and female Long Evans rats trained to self-administer alcohol.

## Materials and Methods

### Animals

Adult male and female Long Evans rats (Envigo-Harlan, Indianapolis, IN) arrived at 7 weeks old and were handled daily for 1 week prior to the start of the experiment. All rats were doubled housed in ventilated cages in same-sex pairs. Rats had ad-libitum access to food and water in the home cage. The rats were kept in a temperature and humidity-controlled colony room that ran on a 12-hour light/dark cycle (lights on at 07:00). All experiments were conducted during the light cycle. Animals were under the care of the veterinary staff from the Division of Comparative Medicine at UNC-Chapel Hill. All of the procedures followed the guidelines established by the NIH Guide to Care and Use of Laboratory Animals and institutional guidelines. All procedures were approved by the UNC-Chapel Hill IACUC. UNC-Chapel Hill is an AAALAC accredited institution.

### Drugs

Ethanol (95% v/v; Pharmco-AAPER, Shelbyville, KY) was diluted in tap water. Poly(I:C) (GE Healthcare and Life Sciences, Pittsburgh, PA; lot #s UF2722 and TB2702) was dissolved in sterile phosphate-buffered saline (PBS), which was also used for control injections. Poly(I:C) was injected intraperitoneal (IP) at a volume of 1 ml/kg.

### Apparatus

Self-administration chambers (Med Associates Inc., St. Albans, VT) were individually located within sound attenuating chambers with an exhaust fan to circulate air and mask outside sounds. Chambers were fitted with a retractable lever on the opposite walls (left and right) of the chamber. There was a cue light above each lever and liquid receptacles in the center panels adjacent both levers. Responses on the left (i.e., active/alcohol) lever resulted in cue light illumination, stimulus tone, and delivery of 0.1 ml of solution across 1.66 seconds via a syringe pump into the left receptacle once the response requirement was met. Responses on the right (inactive) lever had no programmed consequence. The chambers also had infrared photobeams which divided the floor into 4 zones to record general locomotor activity throughout each session.

### EtOH Self-Administration Training

Self-administration sessions (30 minutes) took place 5 days per week (M-F) with the active lever on a fixed ratio 2 schedule of reinforcement such that every second response resulted in delivery of EtOH. A sucrose-fading procedure was used in which EtOH was gradually added to the 10% (w/v) sucrose solution. The exact order of exposure was as follows: 2% (v/v) EtOH/10% (w/v) sucrose (2E/10S), 5E/10S, 10E/10S, 10E/5S, 15E/5S, 15E/2S, 15E, 15E/2S. Following sucrose fading, sweetened EtOH (15E/2S) was the reinforcer for the remainder of the study. At the end of each session, wells were inspected to ensure that rats had consumed all fluid.

#### Experiment 1: Effects of sub-chronic repeated poly(I:C) dosing

To assess the effects of repeated dosing with poly(I:C), rats (n=12 males, 12 females) were injected with poly(I:C) (0 or 0.3 mg/kg, IP, n = 6/group) every other day, 3 hours prior to SA sessions. Standard self-administration sessions continued on non-injection days such that rats received a total of 4 injections and 8 self-administration sessions. Following the 8th SA session, the dose was increased to 1.0 mg/kg for an additional 5 injections with the same alternating-day pattern, then rats were tested in 5 consecutive SA sessions without injection. Finally, on two consecutive days rats were injected with 1.0 mg/kg poly(I:C) 3 hours prior to self-administration sessions on 2 consecutive days and then continued self-administration for 5 more sessions. See figure 1A for design.

**Figure 1.**
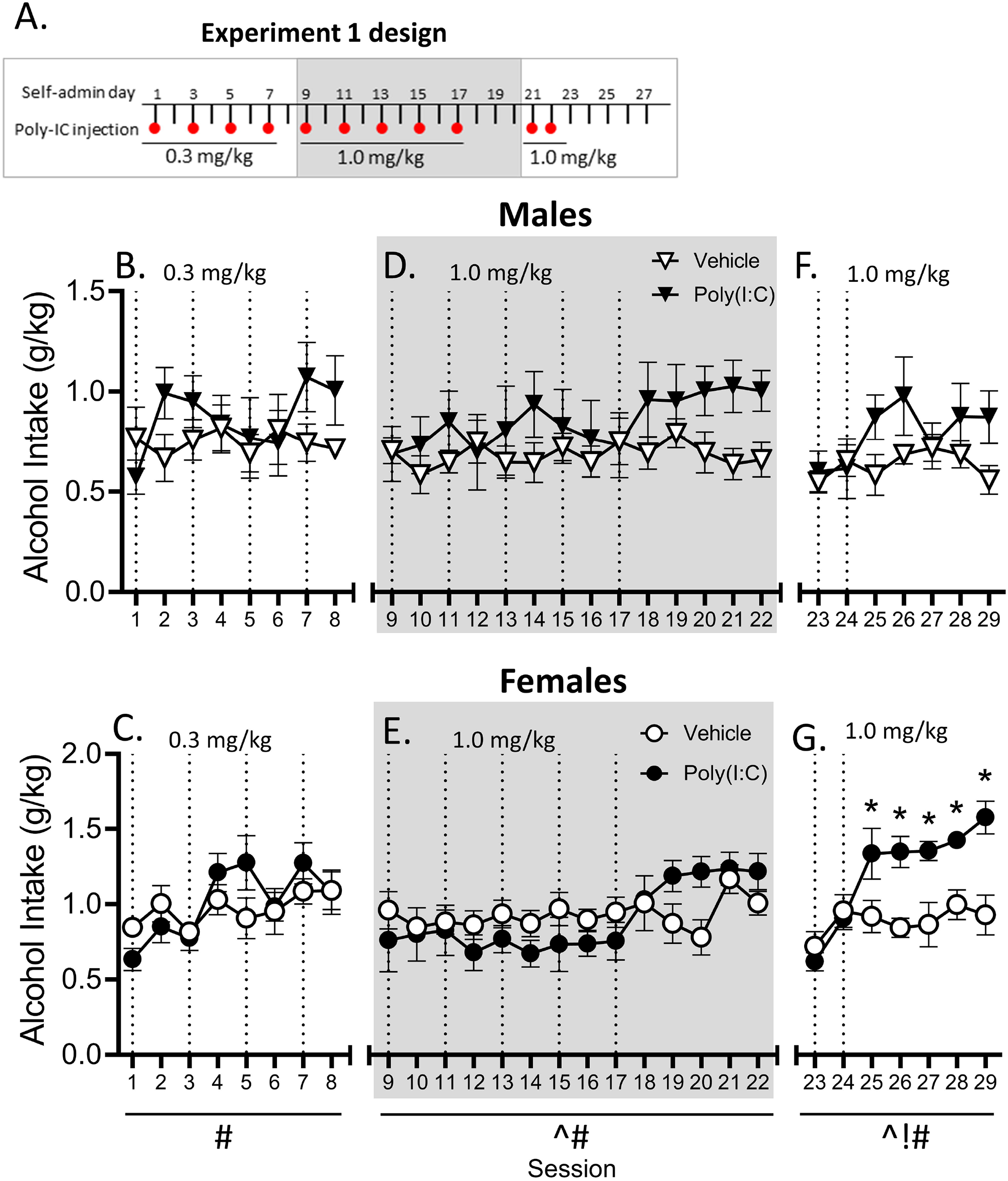
**A**. Experiment 1 design. Male and female rats were trained on daily operant self-administration and were given injections of poly(I:C) 3 hours prior to daily sessions as indicated by dashed lines. **B/C**. A low 0.3 mg/kg dose of poly(I:C) every other day did not affect alcohol drinking. **D/E**. Increasing the dose to 1.0 mg/kg did not affect drinking in males or females, but when two doses were administered consecutively **(F/G)** there was an increase in alcohol consumption in females on the following 5 sessions. ! main effect of poly(I:C), # main effect of session, ^ session x treatment interaction, * p<0.05 vehicle vs. poly(I:C).

#### Experiment 2: Effects of poly(I:C) in naïve rats

To determine whether the effects observed following 2-day administration of poly(I:C) in Experiment 1 were dependent on having a history of poly(I:C) experience, a new group of rats was trained to self-administer alcohol. Following acquisition, rats were injected with poly(I:C) (0 or 1.0 mg/kg, IP, n = 6/group) prior to 2 consecutive SA sessions. Rats continued self-administration sessions for three more days. This protocol was then repeated 2 times for a total of 6 injections over 3 weeks. See figure 3A for design.

#### Experiment 3: Effects of high-dose poly(I:C) on self-administration

In order to determine what effects a higher dose of poly(I:C) would have on alcohol self-administration, a new cohort of male and female rats were trained to self-administer alcohol. Following acquisition, rats were injected with poly(I:C) (0 or 3.0 mg/kg, IP, n = 9-12/group) for 2 consecutive days prior to self-administration sessions and continued self-administration for the remainder of the week. This protocol was repeated twice for a total of 6 injections over 3 weeks. In the following week, the same cohort of rats were injected with a higher dose of poly(I:C) (0 or 9.0 mg/kg, IP, n = 12/group), following the same self-administration protocol, for a single week for a total of 2 injections. This dose was selected based on preliminary studies showing that 9 mg/kg poly(I:C) increased TLR3 gene expression in brain 24 hours post-injection. See figure 4A for experimental design.

### Statistical Analyses

For all experiments, the dependent variables were alcohol lever responses, alcohol intake (g/kg) derived from from the number of reinforcers received and body weight, and locomotor rate (beam breaks/min). All experiments were analyzed using repeated measures analysis of variance (RM-ANOVA) with sex and treatment condition as between-subjects variables. Separate analyses were used for each round of injections (indicated by alternating white and gray sections on figures). Bonferroni was used for all post-hoc analyses.

## Results

### Experiment 1: Sub-chronic injections of poly(I:C) increases alcohol intake in female rats

As shown in figure 1B/C, male and female rats did not show any effects of sub-chronic treatment with poly(I:C) (0.3 mg/kg, IP; figure 1B and C, respectively), while for females there was a general increase in alcohol intake across session (F[7,70]=4.303, p<0.001; figure 1C). When the dose was increased to 1.0 mg/kg every other day (Figures 1D/E), in females there was a main effect of session (F[13,130]=2.85, p<0.01] and a significant interaction (F[13,30]=1.84, p<0.05] while no effects on drinking were seen in males. After rats were injected on two consecutive sessions, females showed an initial increase in alcohol intake (Fig 1G) that lasted for 5 sessions. There was a main effect of session (F[6,60]=9.01, p<0.0001), treatment (F[1,10]=16.44, p<0.01) and a session by treatment interaction in alcohol intake (F[10,60]=2.69, p<0.01). Post-hoc analysis showed that alcohol intake was significantly increased in females on post-injection sessions 1-5 (p’s<0.05). There was also a session by treatment interaction in locomotor rate in females (F[6,60]=2.84, p<0.05). By contrast, males showed no change in alcohol intake and no change in locomotor behavior following consecutive poly(I:C) injections (supplemental figure 2F/G). Results on alcohol lever responses were similar and thus will not be discussed in detail (see table 1 for data and results), and no significant changes were seen in locomotor activity across the experiment (data not shown).

**Figure 2.**
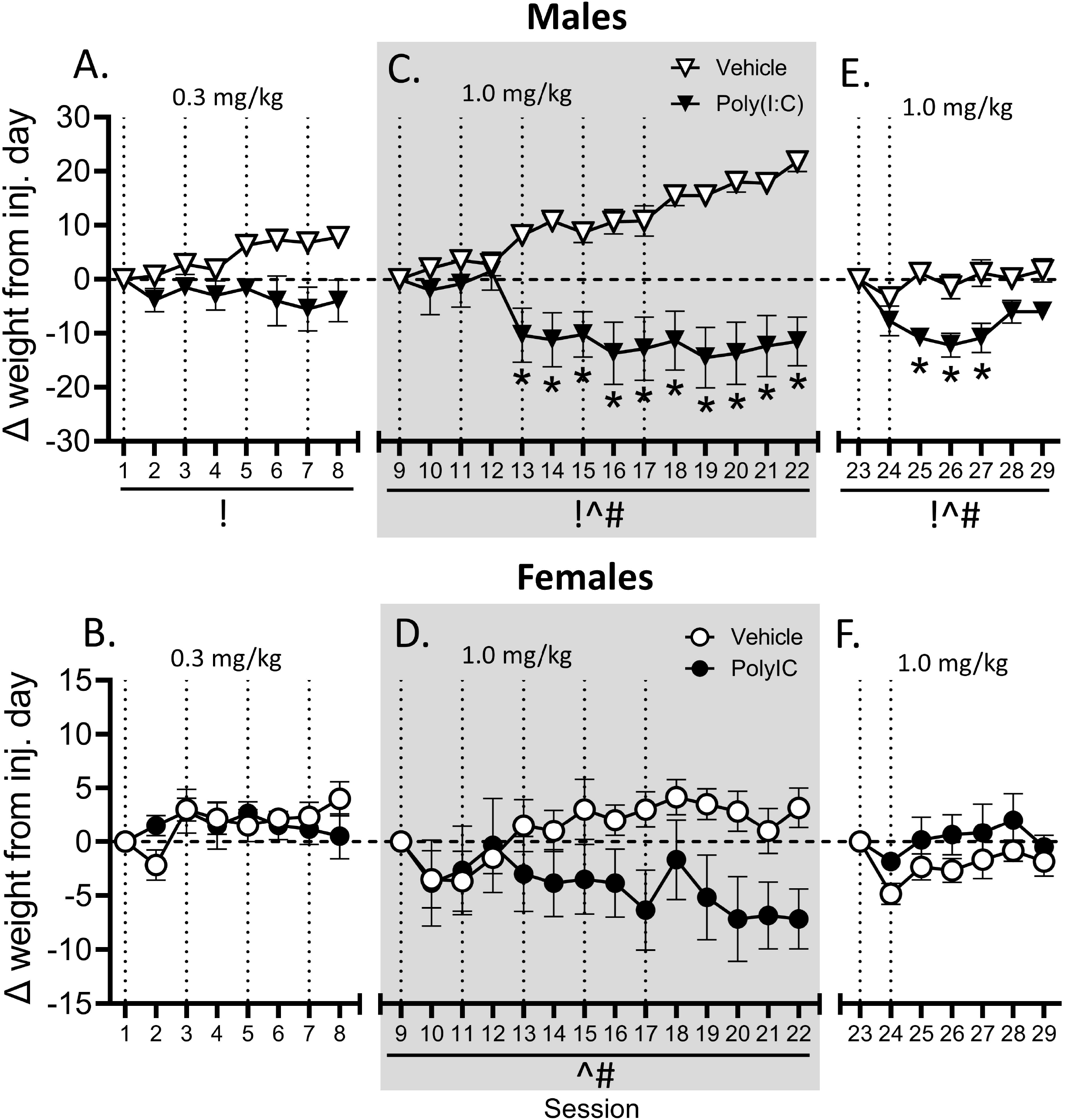
**A-F**. Weight change in experiment 1 relative to the first injection day of each round of injections for males and females. ! main effect of poly(I:C), # main effect of session, ^ session x treatment interaction, * p<0.05 vehicle vs. poly(I:C).

**Table 1.**
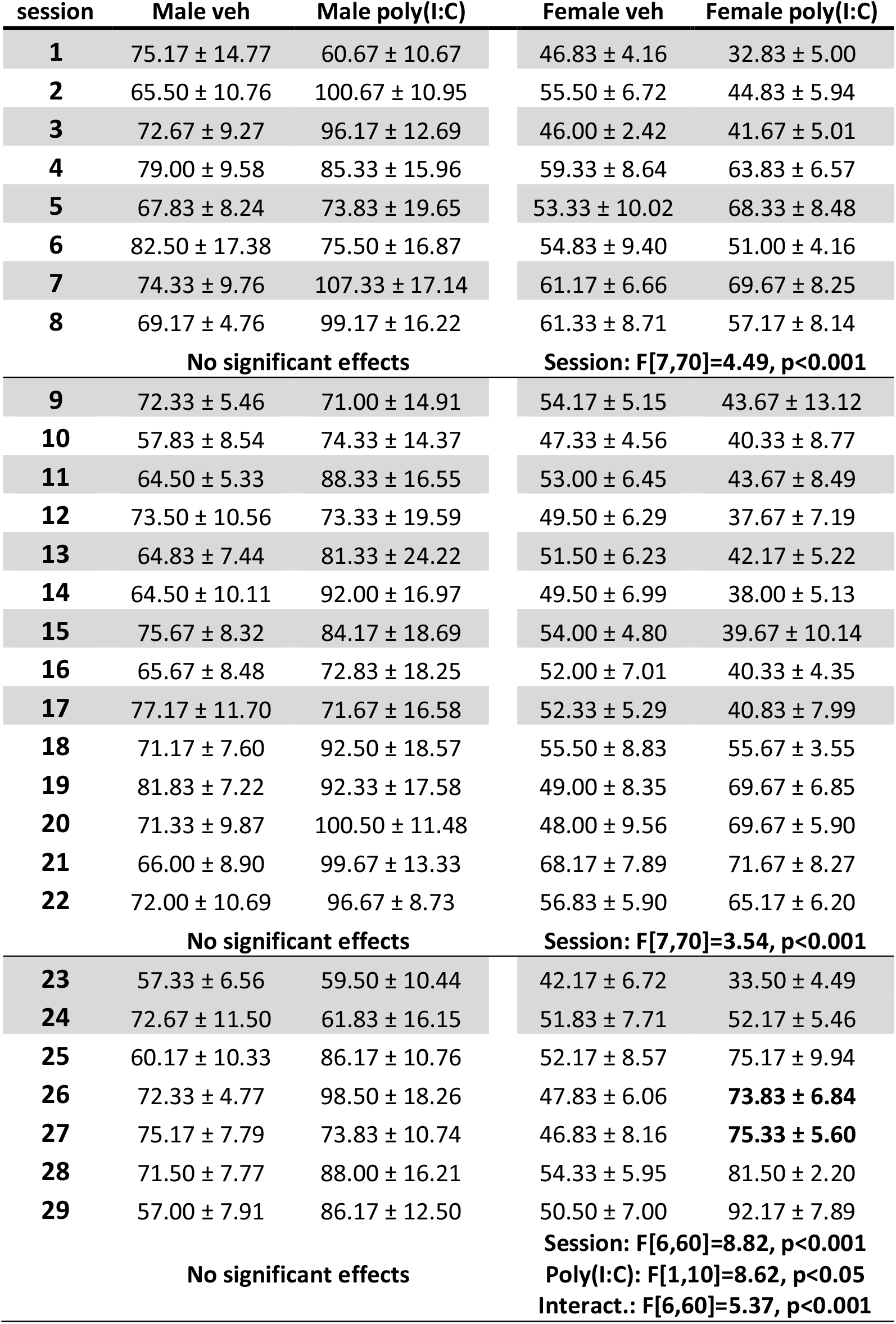
Experiment 1 alcohol lever responses. Number of alcohol lever responses on each day of Experiment 1 (mean ± SEM). Gray indicates imiquimod injection days and horizontal lines delineate phases of the experiment that were analyzed independently. Significant post hoc differences between vehicle and poly(I:C) on each testing day are bolded.

Males that received 0.3 mg/kg poly(I:C) showed a general reduction in weight gain as compared to vehicles (F[1,10]=11.13, p<0.01] while females were unaffected (figure 2A/B). When given injections every other day, in males there were main effects of poly(I:C) (F[1,10]=19.38, p<0.0001), session (F[12,120]=1.98, p<0.05), and a significant interaction (F[12,120]=16.04, p<0.0001) where a difference emerged and was maintained beginning with session 13 (figure 2C). In females there was a main effect of session (F(12,120=1.91, p<0.05) and a significant interaction (F[12,120]=4.32, p<0.0001) with no post hoc differences on any individual day (figure 2D). Lastly, after receiving 1.0 mg/kg on two consecutive days there were main effects of treatment (F[1,10=12.17], p<0.01), session (F[5,50=3.78], p<0.01) and a significant interaction F([5,50]=3.50, p<0.01) where there were differences following the injections on days 25-27 (figure 2E). No differences in weight change were observed in females (figure 2F). Overall males were more sensitive to the poly(I:C), displaying larger and more reliable weight loss.

### Experiment 2: Poly(I:C) treatment enhances alcohol self-administration in female rats

Rats in Experiment 2 were poly(I:C)-naïve. As shown in Figure 3B, there was a main effect of session on alcohol intake in males for the second round of poly(I:C) injections (F[4,40]=3.64, p<0.05) with intake increasing across sessions. In females there was a main effect of poly(I:C) on the third round of injections (F[1,9]=11.99, p<0.01) where poly(I:C) rats drank more than vehicle controls (figure 3C).

While there were no effects of poly(I:C) on alcohol intake in male rats, locomotor rate was affected (figure 3 D). There was a significant interaction on the first round of injections (F[4,40]=5.63, p<0.01) and a reduction in activity on the first injection day (p<0.05). On the second round there was a main effect of day (F[4,40]=3.05, p<0.05) as well as an interaction (F[4,40]=3.61, p<0.05). The same pattern was observed with the third round of injections (main effect: F[3,30]=5.58, p<0.01; interaction: F[3,30]=8.20, p<0.001) with the poly(I:C) injected males showing greater activity two days after the final injection (p<0.05). Results on alcohol lever responses were similar (see table 2). In females no significant change in locomotor activity was found in any of the injection rounds (figure 3E). The pattern of results suggests a male-specific overall enhancement in behavioral activity following poly(I:C) that did not translate to alcohol responding. Lastly, poly(I:C) reduced weight gain in males after the first round of injections (F[3,30=6.61, p<0.01), and there was also a significant interaction (F[3,30=3.56, p<0.05) with a post hoc difference found on the three post-injection days (figure 3F). There was also an effect of session on each of the rounds of injection (first: F[10,30]=7.41,p < 0.0001; second: F[10,30]=6.80, p<0.0001; third: F[10,30]=3.99, p<0.01). Once again in females there were no significant effects found on change in weight (figure 3G).

**Figure 3.**
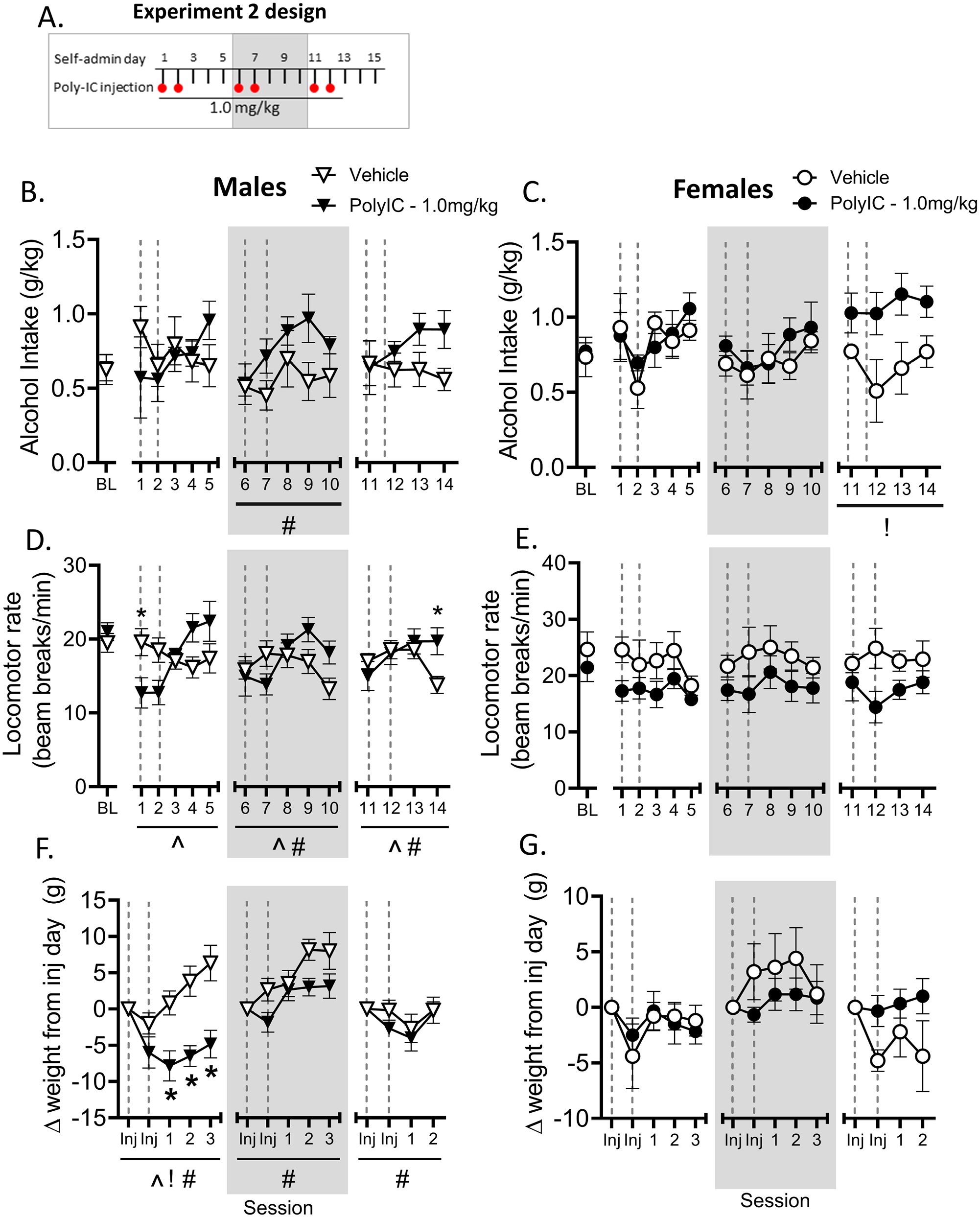
**A**. Experiment 2 design. Male and female rats were trained on daily operant self-administration and were given 1.0 mg/kg injections of poly(I:C) 3 hours prior to daily sessions every other day as indicated by dashed lines. **B/C**. Alcohol intake was not affected in males or females until the third round of injections where poly(I:C) increased alcohol consumption specifically in females. **D**. In males, general locomotor activity was reduced by the very first dose then had little effect until two days after the final dose where rats given poly(I:C) showed greater locomotor activity than controls. **E**. Poly(I:C) did not affect locomotor activity in females. **F/G**. Weight change in experiment 2 relative to the first injection day of each round of injections. ! main effect of poly(I:C), # main effect of session, ^ session x treatment interaction, * p<0.05 vehicle vs. poly(I:C).

**Table 2.**
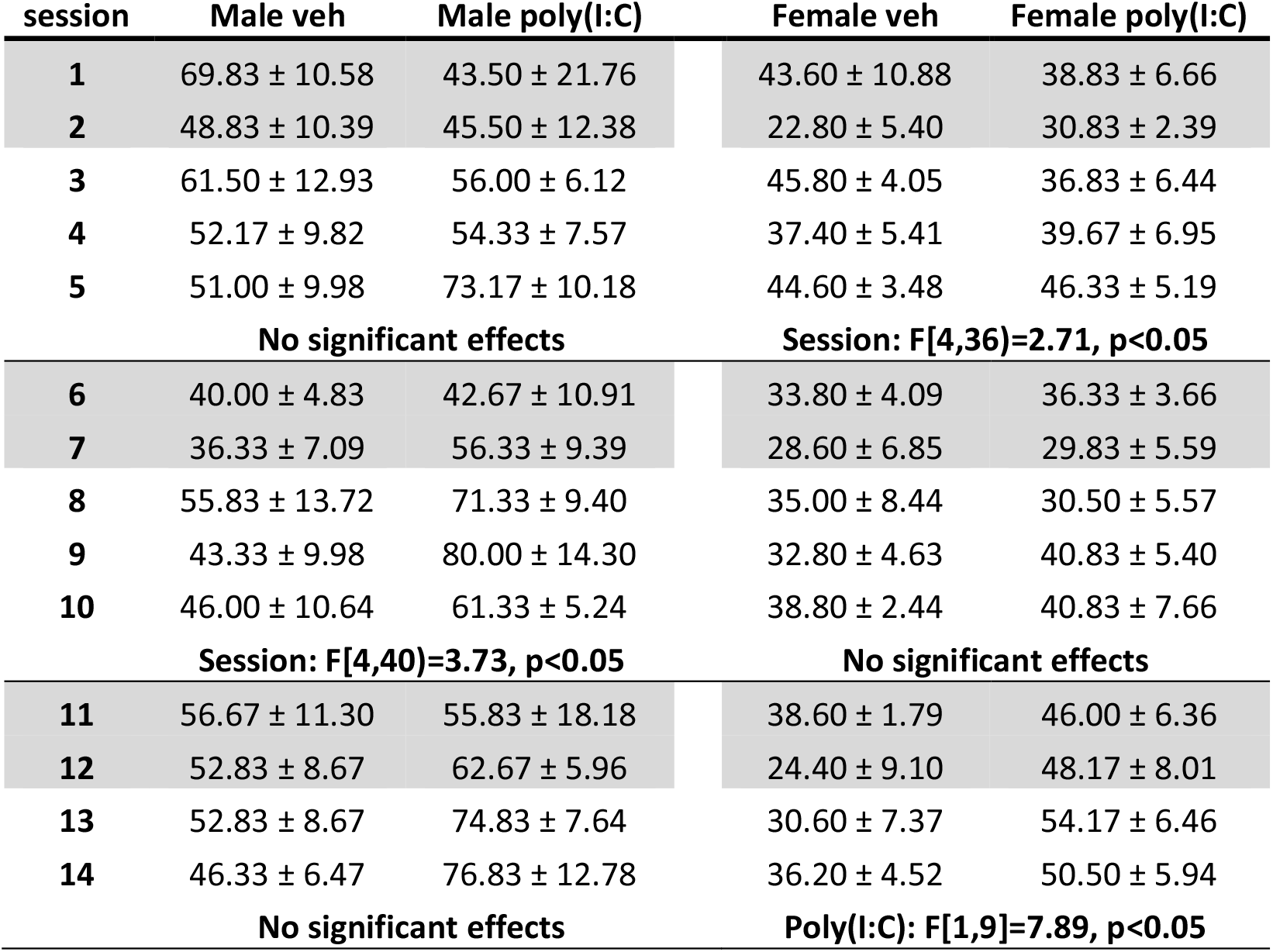
Experiment 2 alcohol lever responses. Number of alcohol lever responses on each day of Experiment 1 (mean ± SEM). Gray indicates imiquimod injection days and horizontal lines delineate phases of the experiment that were analyzed independently.

### Experiment 3: High dose poly(I:C) enhances alcohol self-administration in male rats

As shown in Figure 4B, male rats showed a significantly increased alcohol intake following poly(I:C). There was a main effect of session for the second (at 3.0 mg/kg: F[4,76]=2.83, p<0.05) and fourth rounds of poly(I:C) treatment (at 9.0 mg/kg: F[4,76]=12.80, p<0.001), with alcohol intake increasing across session. Moreover, there was a treatment x session interaction on alcohol intake for the first (F[4,76]=2.80,p<0.05), second (F[4,76]=3.92, p<0.01), and fourth (F[4,76]=12.76, p<0.001)rounds of poly(I:C). Post-hoc analyses found that following 9 mg/kg poly(I:C), male rats had reduced alcohol intake on the first injection day (p<0.05) and consumed more alcohol during the 3^rd^ session post-injection (p<0.05). By contrast, alcohol intake in female rats was reliably decreased on injection days following repeated dosing with poly(I:C). As shown in Figure 4C, there was a main effect of session on alcohol intake for first round (F[4,84]=7.36, p< 0.001), second (F[4,84]=10.43, p<0.001), third (F[4,84]=12.55, p<0.001), and fourth (F[4,84]=10.78, p<0.001) rounds. Moreover, there were a treatment x session interactions on all rounds (first: F[4,84]=6.68, p<0.001); second: F[4,84]=9.49, p<0.001); third: F[4,84]=8.73, p<0.001), and fourth: F[4,84]=4.12, p<0.01). Post-hoc analyses showed that alcohol intake was significantly decreased on the first injection session of each round as well as the second injection of the second round (p’s<0.05), but there were no increases post-injection. Results on alcohol lever responses were similar (see table 3).

**Figure 4.**
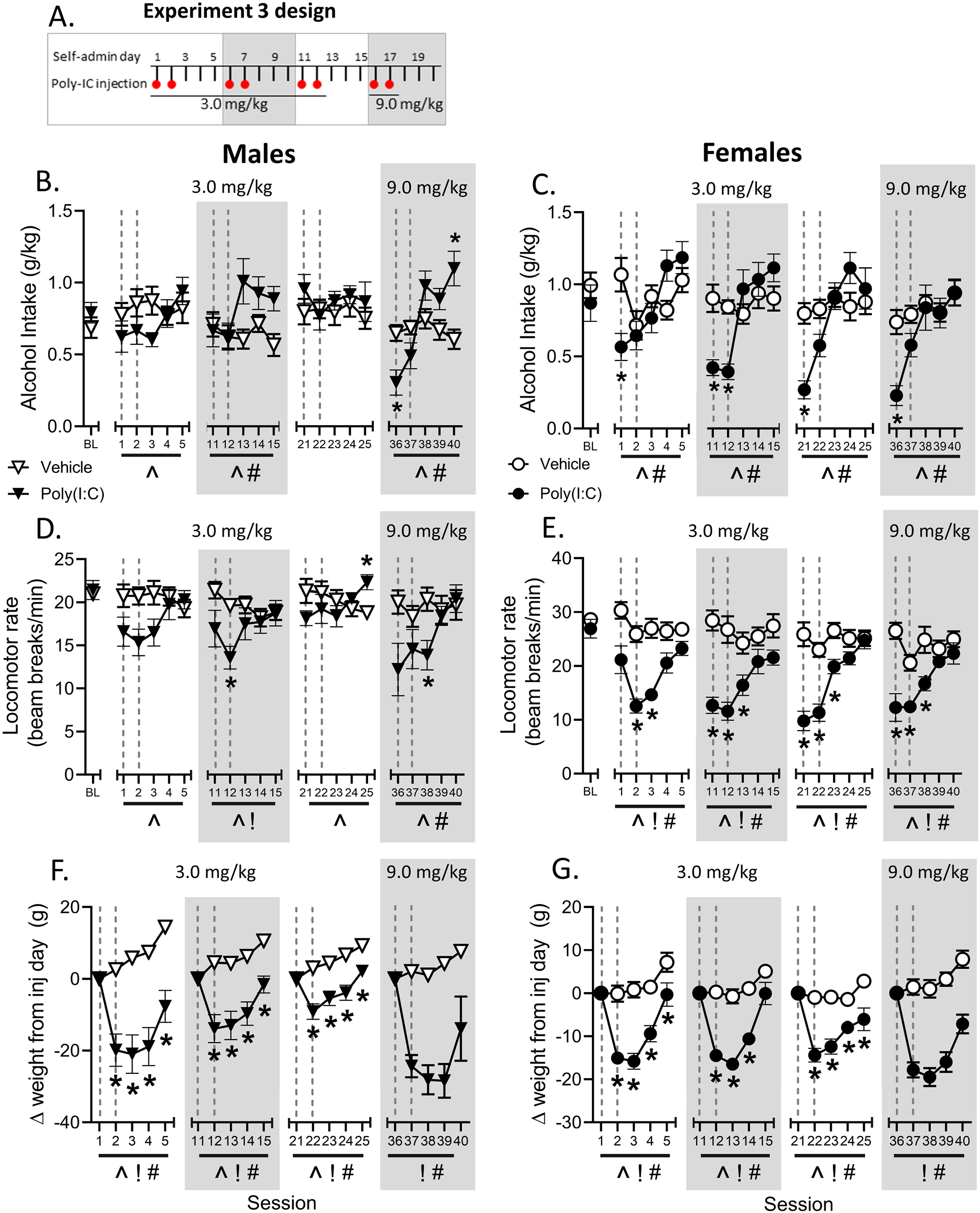
Experiment 3 design. Male and female rats were trained on daily operant self-administration and were given 3.0 mg/kg injections of poly(I:C) 3 hours prior to daily sessions every other day for three rounds of injections, then finally were given injections of 9.0 mg/kg on two subsequent days. **B/C**. Males primarily showed changes in alcohol intake at the highest dose whereas females reliably drank less alcohol on the first injection day. **D/E**. Locomotor activity in both sexes was generally reduced by poly:IC but it was particularly strong in females. **F/G**. Weight change in experiment 3 relative to the first injection day of each round of injections.! main effect of poly(I:C), # main effect of session, ^ session x treatment interaction, * p<0.05 vehicle vs. poly(I:C).

**Table 3.**
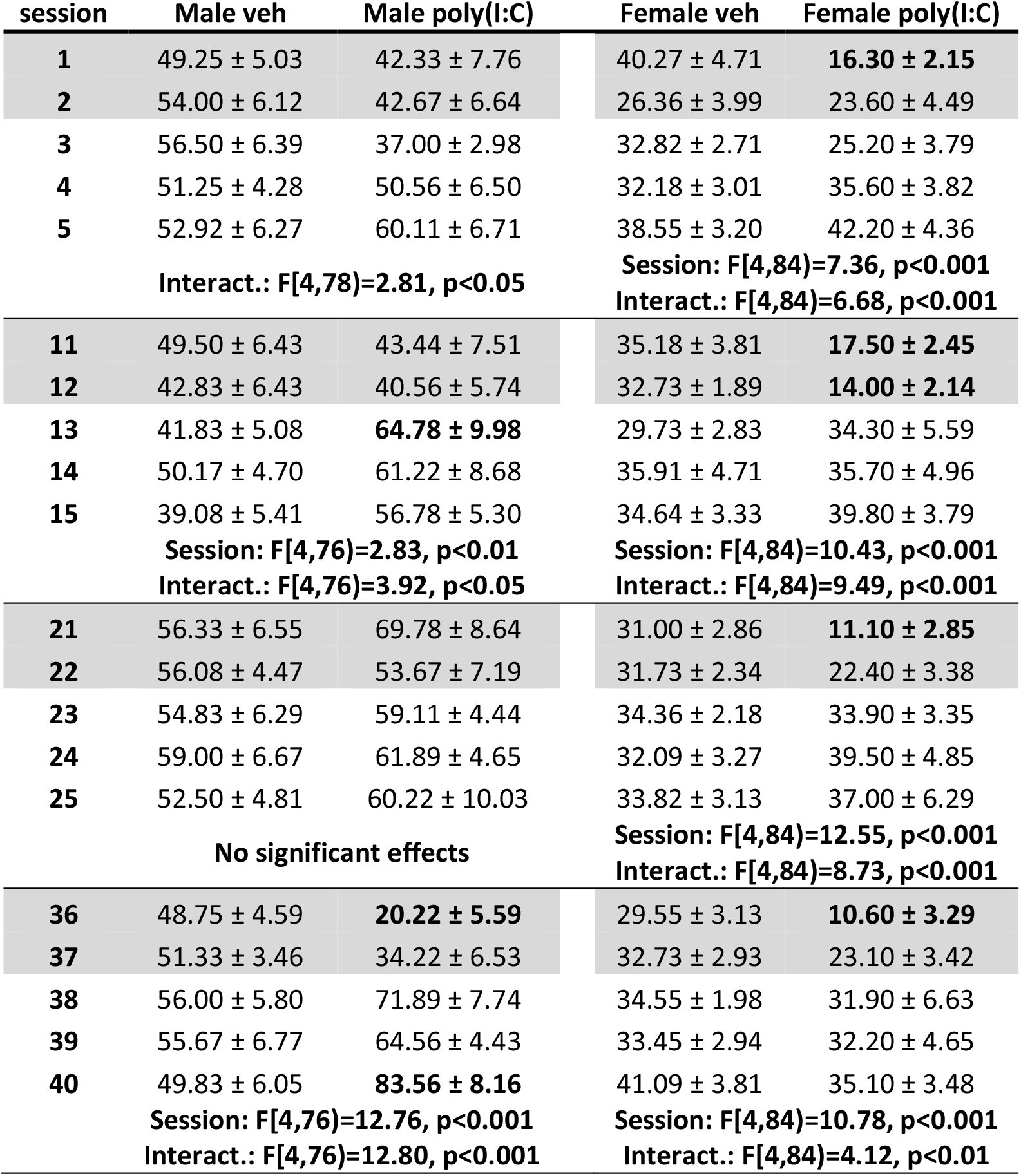
Experiment 3 alcohol lever responses. Number of alcohol lever responses on each day of Experiment 1 (mean ± SEM). Gray indicates imiquimod injection days and horizontal lines delineate phases of the experiment that were analyzed independently. Significant post hoc differences between vehicle and poly(I:C) on each testing day are bolded.

| session | Male veh | Male poly(I:C) | Female veh | Female poly(I:C) |
| --- | --- | --- | --- | --- |
| **1** | 49.25 ± 5.03 | 42.33 ± 7.76 | 40.27 ± 4.71 | **16.30 ± 2.15** |
| **2** | 54.00 ± 6.12 | 42.67 ± 6.64 | 26.36 ± 3.99 | 23.60 ± 4.49 |
| **3** | 56.50 ± 6.39 | 37.00 ± 2.98 | 32.82 ± 2.71 | 25.20 ± 3.79 |
| **4** | 51.25 ± 4.28 | 50.56 ± 6.50 | 32.18 ± 3.01 | 35.60 ± 3.82 |
| **5** | 52.92 ± 6.27 | 60.11 ± 6.71 | 38.55 ± 3.20 | 42.20 ± 4.36 |
|  | **Interact.: F[4,78]=2.81, p<0.05** |  | **Session: F[4,84]=7.36, p<0.001**<br>**Interact.: F[4,84]=6.68, p<0.001** |  |
| **11** | 49.50 ± 6.43 | 43.44 ± 7.51 | 35.18 ± 3.81 | **17.50 ± 2.45** |
| **12** | 42.83 ± 6.43 | 40.56 ± 5.74 | 32.73 ± 1.89 | **14.00 ± 2.14** |
| **13** | 41.83 ± 5.08 | **64.78 ± 9.98** | 29.73 ± 2.83 | 34.30 ± 5.59 |
| **14** | 50.17 ± 4.70 | 61.22 ± 8.68 | 35.91 ± 4.71 | 35.70 ± 4.96 |
| **15** | 39.08 ± 5.41 | 56.78 ± 5.30 | 34.64 ± 3.33 | 39.80 ± 3.79 |
|  | **Session: F[4,76]=2.83, p<0.01**<br>**Interact.: F[4,76]=3.92, p<0.05** |  | **Session: F[4,84]=10.43, p<0.001**<br>**Interact.: F[4,84]=9.49, p<0.001** |  |
| **21** | 56.33 ± 6.55 | 69.78 ± 8.64 | 31.00 ± 2.86 | **11.10 ± 2.85** |
| **22** | 56.08 ± 4.47 | 53.67 ± 7.19 | 31.73 ± 2.34 | 22.40 ± 3.38 |
| **23** | 54.83 ± 6.29 | 59.11 ± 4.44 | 34.36 ± 2.18 | 33.90 ± 3.35 |
| **24** | 59.00 ± 6.67 | 61.89 ± 4.65 | 32.09 ± 3.27 | 39.50 ± 4.85 |
| **25** | 52.50 ± 4.81 | 60.22 ± 10.03 | 33.82 ± 3.13 | 37.00 ± 6.29 |
|  | **No significant effects** |  | **Session: F[4,84]=12.55, p<0.001**<br>**Interact.: F[4,84]=8.73, p<0.001** |  |
| **36** | 48.75 ± 4.59 | **20.22 ± 5.59** | 29.55 ± 3.13 | **10.60 ± 3.29** |
| **37** | 51.33 ± 3.46 | 34.22 ± 6.53 | 32.73 ± 2.93 | 23.10 ± 3.42 |
| **38** | 56.00 ± 5.80 | 71.89 ± 7.74 | 34.55 ± 1.98 | 31.90 ± 6.63 |
| **39** | 55.67 ± 6.77 | 64.56 ± 4.43 | 33.45 ± 2.94 | 32.20 ± 4.65 |
| **40** | 49.83 ± 6.05 | **83.56 ± 8.16** | 41.09 ± 3.81 | 35.10 ± 3.48 |
|  | **Session: F[4,76]=12.76, p<0.001**<br>**Interact.: F[4,76]=12.80, p<0.001** |  | **Session: F[4,84]=10.78, p<0.001**<br>**Interact.: F[4,84]=4.12, p<0.01** |  |

Interestingly, despite differing effects on alcohol intake, both males and females showed decreases in locomotor rate following poly(I:C) treatment. As shown in Figure 4D, in males there was a session by treatment interaction on locomotor rate for the first round (F[4,76]=3.98, p<0.01), second round (F[4,76]=3.59, p<0.01), third round (F[4,76]=4.96, p<0.001), and fourth round (F[4,76]=5.09, p<0.001). Post hoc analyses showed that locomotor rate was decreased following the second injection of round 2 and again on the third session of round 4 (p’s<0.05), as well as an increase on the final session of round 3 (p<0.05), suggesting mild overall motor effects in males. In contrast, females showed more marked effects on locomotor rate (figure 4E). There was a main effect of session for each round (first: F[4,66]=9.95, p<0.001; second: F[4,66]=6.45, p<0.001; third: F[4,66]=19.09, p<0.0001; fourth: F[4,66]=8.45, p<0.001) as well as a session by treatment interaction for each round (first: F[4,84]=4.70, p<0.001; second: F[4,84]=6.45, p<0.001; third: F[4,84]=14.34, p<0.001; fourth: F[4,84]=8.45, p<0.001). Finally, there was also a main effect of treatment on locomotor rate in females for all rounds (first: F[1,21]=28.81, p<0.001; second: F[1,21]=30.22, p<0.001; third: F[1,21]=28.18, p<0.001; fourth: F[1,21]=22.88, p<0.001). Post hoc analysis found that locomotor rate was significantly decreased in the poly(I:C) females on every injection day (except the first) as well as day after the second injection on each round (p’s<0.05; figure 3E). Taken together, these effects suggest sex differences in response to poly(I:C) where males show increases in drinking after repeated poly(I:C) injections and females are sensitive to poly(I:C) on injection days.

Lastly, there were robust differences in weight change after every injection in both males and females (figure 4F/G). Across all injections there were main effects of poly(I:C) injection (males round 1: F[1,19]=30.07, p<0.0001); round 2: F[1,19]=27.46, p<0.0001); round 3: F[1,19]=21.18, p<0.001; round 4: F[1,19]=56.82, p<0.0001); females round 1: F[1,21]=39.45, p<0.0001); round 2: F[1,21]=28.10, p<0.0001); round 3: F[1,21]=31.50, p<0.0001), round 4: F[1,21]=53.95, p<0.0001) and session (males round 1: F[3,57]=68.39, p<0.0001); round 2: F[3,57]=48.02, p<0.0001); round 3: F[3,57]=56.13, p<0.001; round 4: F[3,57]=5.08, p<0.01); females round 1: F[3,63]=39.45, p<0.0001); round 2: F[3,63]=39.75, p<0.0001), round 3: F[3,63]=9.71, p<0.0001); round 4: F[3,63]=33.22, p<0.0001). There were also interactions for all three rounds of 3.0 mg/kg injections in males (round 1: F[3,57]=3.53, p<0.05); round 2: F[3,57]=4.94, p<0.01); round 3: F[3,57]=4.62, p<0.01) and females (round 1: F[3,63]=6.26, p<0.001); round 2: F[3,63]=9.35, p<0.0001); round 3: F[3,63]=3.07, p<0.05) but no interaction when the poly(I:C) dose was increased to 9.0 mg/kg in either sex.

## Discussion

There is considerable evidence that the neuroimmune system plays an important role in alcohol use and the development and maintenance of alcohol use disorder (Coleman and Crews, 2018; Crews et al., 2015; Cui et al., 2014; Erickson et al., 2019; Mayfield et al., 2013). The current experiments add to this body of evidence demonstrating that activation of TLR3 increases alcohol self-administration. Moreover, this work adds to a growing body of literature identifying differential response to the consequences of TLR3 activation on alcohol intake in both males and females (Warden et al., 2019a, b).

First, we found that dosing with poly(I:C) every other day with 0.3 mg/kg was insufficient to increase alcohol intake (figure 1B/C). When the dose was increased to 1.0 mg/kg there was a significant interaction in females but no effect of poly(I:C) (figure 1D/E). To mitigate the potential issues presented by every-other-day dosing, the same rats were injected with 1 mg/kg poly(I:C) on two consecutive days and increased alcohol self-administration was observed for five consecutive days in female rats (Figure 1G). A potential explanation for this finding is that by this point in testing, the neuroimmune system was sufficiently primed by prior poly(I:C) experience such that the reinforcing effects of alcohol were enhanced. Previous studies of innate immune activation have shown that events that continually activate the neuroimmune system sensitize the inflammatory response, creating a phenotype that is predisposed to substance use (Frank et al., 2011). Rats in the current experiments had received 9 injections of poly(I:C) by this point and thus repeated activation of these neuroimmune cascades may have led to sensitization in the system whereby alcohol intake was enhanced. Interestingly, the two consecutive day dosing protocol had no effect on self-administration in the males, again suggesting a differential response to the consequences of poly(I:C) on alcohol self-administration between the sexes. It is also possible that dosing with poly(I:C) every other day may be too frequent, leading to malaise that would otherwise translate to greater alcohol intake if there was no work requirement. Indeed, there were significant weight reductions in males following treatment with both doses and in the females at the 1.0 mg/kg dose (figure 2) that are likely representative of sickness behavior (Careaga et al., 2018), indicating a sex difference in the sensitivity to the poly(I:C)-induced weight loss.

Given that rats in Experiment 1 had extensive experience with poly(I:C) and that the two-day dosing pattern produced a greater increase in drinking in females, in Experiment 2 we sought to determine whether the effects of the two-day dosing paradigm were the result of back-to-back dosing or were due to sensitization after repeated poly(I:C) injections over time. To get at this question, the goal of Experiment 2 was to test whether self-administration would be increased following two consecutive days of poly(I:C) treatment in rats without a poly(I:C) history. We found that the same 1 mg/kg two-day dosing protocol used in Experiment 1 in poly(I:C)-naïve rats did not alter self-administration in males or females on the first cycle (figure 3B/C). Thus, it is indeed likely that repeated immune activation is necessary to observe escalations in alcohol self-administration. Further, the first round of injections reduced both locomotor activity (figure 2D) and loss of weight (figure 3F/G) in males but not females, indicating a sex difference in behavioral sensitivity to poly(I:C). Interestingly, on the injection third cycle, an overall escalation in self-administration was found in the poly(I:C)-treated females but not in males (figure 2C). Thus, as was seen in experiment 1 (figure 1F/G) we once again found that females show escalations in drinking after a two-day poly(I:C) challenge, but only after a history of poly(I:C) administration.

Finally, using the same two-day injection protocol cycle but a higher poly(I:C) dose (3 mg/kg), the injections did not produce consistent increases in self-administration (figure 4 B/C). On injection days females, but not males, showed decreases in self-administration. This decrease in self-administration was likely during peak neuroimmune activation which has previously been shown to reduce alcohol intake in female mice (Warden et al., 2019b). It may also be related to an overall motor impairment or sickness behavior as a decrease in locomotor rate was also observed on the injection days in the females (figure 4E). Locomotor behavior was also consistently reduced in females the day after the second injection of each cycle, but self-administration was not altered, which suggests that the rats overcame the motor reduction to self-administer alcohol. In males there was a significant interaction on the second round of injections but no post-hoc tests were significant, thus they were largely unaffected until the dose was increased as increased to 9 mg/kg where the first injection day reduced intake. Subsequently, on the last day of testing they consumed more alcohol (figure 4B). Notably, females did not show this increase at the higher dose. Both the 3.0 and 9.0 mg/kg doses reliably induced weight loss in both males and females, indicative of a maintenance of immune responding and sickness responses. We previously saw similarly present but inconsistent effects on self-administration and locomotor activity alongside consistent weight loss with injection of the TLR7 agonist imiquimod (Lovelock et al. 2021, *submitted*), which may be affecting behaviors through similar signaling pathways as poly(I:C). While the poly(I:C) is clearly having effects, it appears that females are more behaviorally sensitive to poly(I:C) 3 hours post-injection showing locomotor reductions on injection days at lower doses than males. It may be that increasing the dose in males is able to produce a similar behavioral pattern that was seen in females.

An important consideration for the current findings is that the time course of neuroimmune-related changes may differ between males and females. Indeed, previous studies show that peak neuroimmune activation following poly(I:C) treatment differs by up to 24 hours between males and females (Warden et al., 2019a, b). Moreover, given the extensive poly(I:C) exposure that rats in Experiments 1 and 3 had, results may reflect the differential effects of neuroimmune priming between males and females. It is well-established that exposure to stressors can potentiate subsequent neuroimmune and microglial responses to immune challenges (Frank et al., 2011; Johnson et al., 2004; Johnson et al., 2003; Munhoz et al., 2006; Wohleb et al., 2011), and several studies identified that while both males and females are affected by priming of the neuroimmune response, the locus of that effect differs (Bekhbat et al., 2019; Bekhbat et al., 2021; Fonken et al., 2018; Gildawie et al., 2020). Priming of microglia (Melbourne et al., 2021; Walter et al., 2017) and specifically TLR responses (Crews and Vetreno, 2018) are thought to play an important role in AUD, thus a history of immune activation, whether through stress, pathogen exposure, or sterile central neuroimmune activation, may contribute to the onset and/or sustaining of AUD. Considered in this context, the current studies repeatedly applied the same immune challenge rather than inducing priming via stress or other insults, but repeated activation of TLRs appears to result in enhanced responding that may be due to adaptations similar to those seen with stress-induced priming.

The current studies reinforce the importance of the TLR3 signaling pathway in alcohol use. Similar to our previous work and others, activation of TLR3 enhanced alcohol intake (Randall et al., 2019; Warden et al., 2019a, b). Inhibition of the TLR3 signaling pathway via a TLR3 inhibitor decreases alcohol intake in mice (Warden et al., 2019a) while selective inhibition of the TRIF adapter protein via the IKKI and TBK1 components has been shown to decrease alcohol intake in mice (McCarthy et al., 2018), demonstrating a crucial role for both TLR3 and the TRIF pathway in excessive alcohol consumption. Given the complexity of the TRIF adapter system, future studies in rats should assess the contribution of its component parts – IKKI and TBK1. Moreover, an in-depth assessment of both individual differences in neuroimmune response to poly(I:C) and an assessment of other strains beyond Long-Evans to determine whether there exist strain differences in response to poly(I:C) in rats. Finally, greater focus on the timecourse of neuroimmune activation as well as exposure frequency and distribution will be important for understanding how these factors drive alcohol consumption. Overall, the current findings add to the body of work demonstrating the importance of the neuroimmune system in alcohol use.

## Notes

**Funding:** This work was supported in part by the National Institute of Health (AA011605, AA026537) and by the Bowles Center for Alcohol Studies.

### Competing Interest Statement

The authors have declared no competing interest.

